# Social information impairs reward learning in depressive subjects: behavioral and computational characterization

**DOI:** 10.1101/378281

**Authors:** Lou Safra, Coralie Chevallier, Stefano Palminteri

## Abstract

Depression is characterized by a marked decrease in social interactions and blunted sensitivity to rewards. Surprisingly, despite the importance of social deficits in depression, non-social aspects have been disproportionally investigated. As a consequence, the cognitive mechanisms underlying atypical decision-making in social contexts in depression are poorly understood. In the present study, we investigate whether deficits in reward processing interact with the social context and how this interaction is affected by self-reported depression and anxiety symptoms. Two cohorts of subjects (discovery and replication sample: *N* = 50 each) took part in a task involving reward learning in a social context with different levels of social information (absent, partial and complete). Behavioral analyses revealed a specific detrimental effect of depressive symptoms – but not anxiety – on behavioral performance in the presence of social information, i.e. when participants were informed about the choices of another player. Model-based analyses further characterized the computational nature of this deficit as a negative audience effect, rather than a deficit in the way others’ choices and rewards are integrated in decision making. To conclude, our results shed light on the cognitive and computational mechanisms underlying the interaction between social cognition, reward learning and decision-making in depressive disorders.

## Introduction

One of the core clinical symptoms of depression is anhedonia, which refers to a reduced motivation to engage in daily life activities (motivational anhedonia) and a reduced enjoyment of usually enjoyable activities (consummatory anhedonia) *(1, 2)*. In principle, this clinical manifestation could be explained by reduced reward sensitivity, both in terms of incentive motivation and in terms of reinforcement processes *(3–5).* A direct prediction of this hypothesis is that depressive symptoms should be associated with reduced reward sensitivity in learning contexts both at the behavioral and neural level. However, while some studies do find evidence that depressive symptoms in the general population and in clinical depression are associated with blunted reward learning and reward-related signals in the brain *(6, 7)*, others indicate no *(8, 9)* or mixed effects *(5)*. As a consequence, there is no strong consensus about which components of reward processing are most predictive of depressive symptoms in both the general population and clinical depression *(5)*.

Another striking clinical manifestation of depressed symptoms is a marked decrease in social interactions. Depression is indeed associated with social risk factors, social impairments and poor social functioning *(10).* Surprisingly, despite the importance of the socio-cognitive impairments that are often associated with elevated depressive symptoms, non-social aspects have received disproportionate attention. Furthermore, when social aspects are investigated the focus is often on emotional processing and theory of mind but not on how social information is integrated to produce efficient goal-directed behavior *(11)*. In the present study, our goal was to investigate whether the reward-learning deficit that is often associated with elevated depressive symptoms interacts with the social context *(12)*.

According to social learning theory, a sizable amount of decisions are not directly shaped by people’s personal history of reward and punishments, but are rather acquired through social observation *(13)*. More specifically, this framework posits that human learning occurs mostly in social contexts, where subjects can be influenced by social cues (i.e. others’ choices and outcomes) *(13, 14)*. In order to test how depressive symptoms affect the integration of social cues during reinforcement learning, we administered a variant of a previously validated observational learning task on two independent samples of participants: an exploration sample and a replication sample *(14, 15)*. Subjects also completed psychometric questionnaires assessing depression and anxiety (a co-morbid trait). The task included a ‘Private’ learning condition, in which participants only had access to the outcome of their own choice, and two social conditions: the ‘Social-Choice’ condition in which participants had access to the demonstrator’s choice, and the ‘Social-Choice+Outcome’ condition in which participants had access to the demonstrator’s actions and their outcome (**Figure 1**).

**Figure 1:**
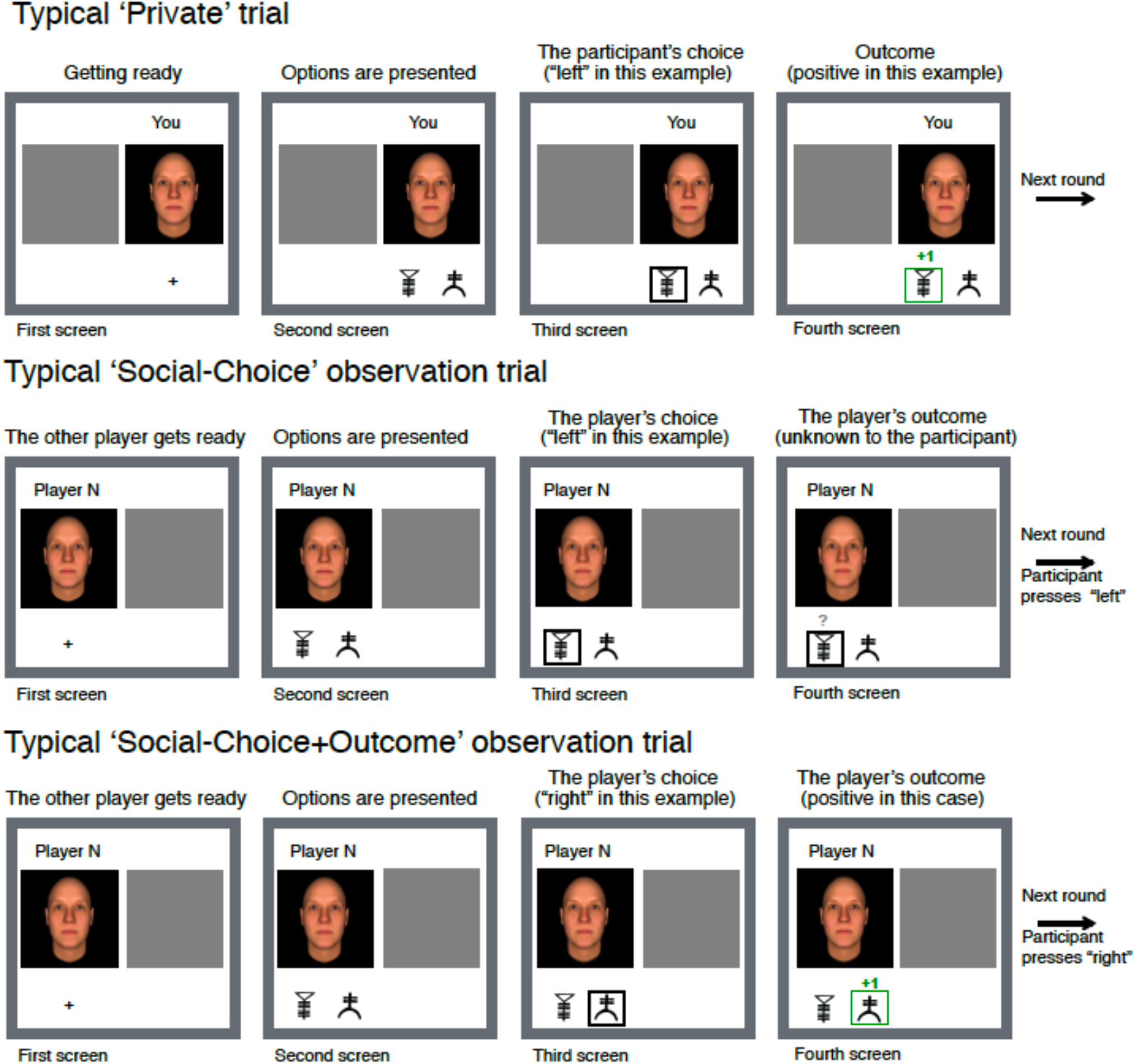
behavioral task. In each condition, participants played in turn with a simulated demonstrator. In each private trial, after each choice, participants received a reward or a punishment. In the Private condition, participants did not see the choice or the outcome of the demonstrator. In the Choice observation condition, the choice of their demonstrator was displayed at each trial. In the Social-Choice+Outcome condition, both the choice and the outcome of the demonstrator were displayed. Note that the Social-Choice and the Social-Choice+Outcome also involved private trials.

Our design allowed us to test several hypotheses concerning the relation between depressive symptoms and learning performance in private and social contexts. First, our design allowed us to test whether or not depressive symptoms degrade reward learning *per se*, as assumed by the standard account of depression as a reward sensitivity deficit. Second, by comparing the ‘Private’ and the ‘Social’ learning contexts, we can assess whether or not depressive symptoms are associated to a learning deficit in ‘Social’ contexts, as predicted by evidence of socio-cognitive impairments in depressive patients. Finally, thanks to computational analyses, we can precisely characterize the learning deficit in the ‘Social’ context either as a primary social learning deficit (i.e., impaired imitation) or as a secondary social learning (i.e., a negative audience effect).

## Results

### Experimental protocol and quality checks

An online experiment was particularly suited to test our hypothesis because - compared to laboratory-based experiments - it provides a more diversified pool of subjects, in terms of psychiatric traits and cognitive performance *(16–19)*. Specifically, we tested 50 participants in the general population and then ran a direct replication of the experiment on a second independent sample of 50 participants. Levels of depressive and anxiety symptoms were assessed and spanned a large range (**Table 1**) *(20)*, with good internal consistency (Hospital Anxiety Depression scale - depression subscale: Cronbach’s alpha 85%; anxiety subscale: Cronbach’s alpha 84%).

**Table 1:** descriptive statistics for age, gender, depression and anxiety scores. For each sample, the mean of each demographic variable is presented with its 95% confidence interval.

|  | Age | Sex ratio<br>(% women) | Depression | Anxiety |
| --- | --- | --- | --- | --- |
| First sample<br>(N = 50) | 33.02 ± 1.25<br>[22 – 62] | 28% | 5.46 ± 1.26<br>[0 – 19] | 6.40 ± 1.16<br>[0 – 15] |
| Second sample<br>(N = 50) | 33.76 ± 3.28<br>[19 – 61] | 42% | 4.96 ± 1.27<br>[0 – 16] | 6.30 ± 1.28<br>[0 – 20] |
| Statistical<br>difference | t(98) = 0.36<br>p > .250 | X-squared = 1.58,<br>df = 1, p-value = .208 | t(98) = 0.56 p > .250 | t(98) = 0.12 p > .250 |

Participants were paired with a virtual demonstrator and performed a probabilistic reinforcement learning task in three contexts: a ‘Private’, in which participants performed the task individually with no access to the demonstrator’s choices and outcomes, and two social conditions: the ‘Social-Choice’ condition in which participants had access to the demonstrator’s choices, and the ‘Social-Choice+Outcome’ condition in which participants had access to the demonstrator’s choices and their outcome. Overall, participants displayed robust instrumental learning by choosing the most rewarded symbol above chance in all conditions (‘Private’: *M* = 0.65 ± 0.03, *t*(99) = 11.42, *p* < .001; ‘Social-Choice’: *M* = 0.65 ± 0.03, *t*(99) = 11.63, *p* < .001; ‘Social-Choice+Outcome’: *M* = 0.67 ± 0.03, *t*(99) = 12.36, *p* < .001; ± corresponds to the 95% confidence intervals; Figure 1A).

### Assessing observational learning

Contrary to previous studies *(14, 15)*, we used an online adaptive learning algorithm that determined the demonstrator’s behavior (Q-learning with learning rate = 0.5 and choice temperature = 10). As a consequence, the virtual demonstrators displayed realistic learning curves with some variability of performance. We predicted that observational learning would result in a correlation between the correct choice rate of the participants and that of the demonstrator in a given learning session. To test this prediction, we used a mixed linear regression with ‘Condition’ (‘Private’ vs ‘Social-Choice’ vs ‘Social-Choice+Outcome’) as a within-subject factor and the demonstrator’s performance in a given learning session as a between-subject variable. As predicted, a higher demonstrator’s percentage of correct choices (i.e., ‘good’ demonstrations) was associated with a higher participants’ rates of correct choices in both social conditions (‘Social-Choice’ *vs* ‘Private’: *t*(495) = 2.70, *p* = .007; ‘Social-Choice+Outcome’ *vs* ‘Private’: *t*(495) = 2.25, *p* =.025) but not in the Private condition (t(495) = 0.10, *p* > .250; **Figure 2A**).

**Figure 2:**
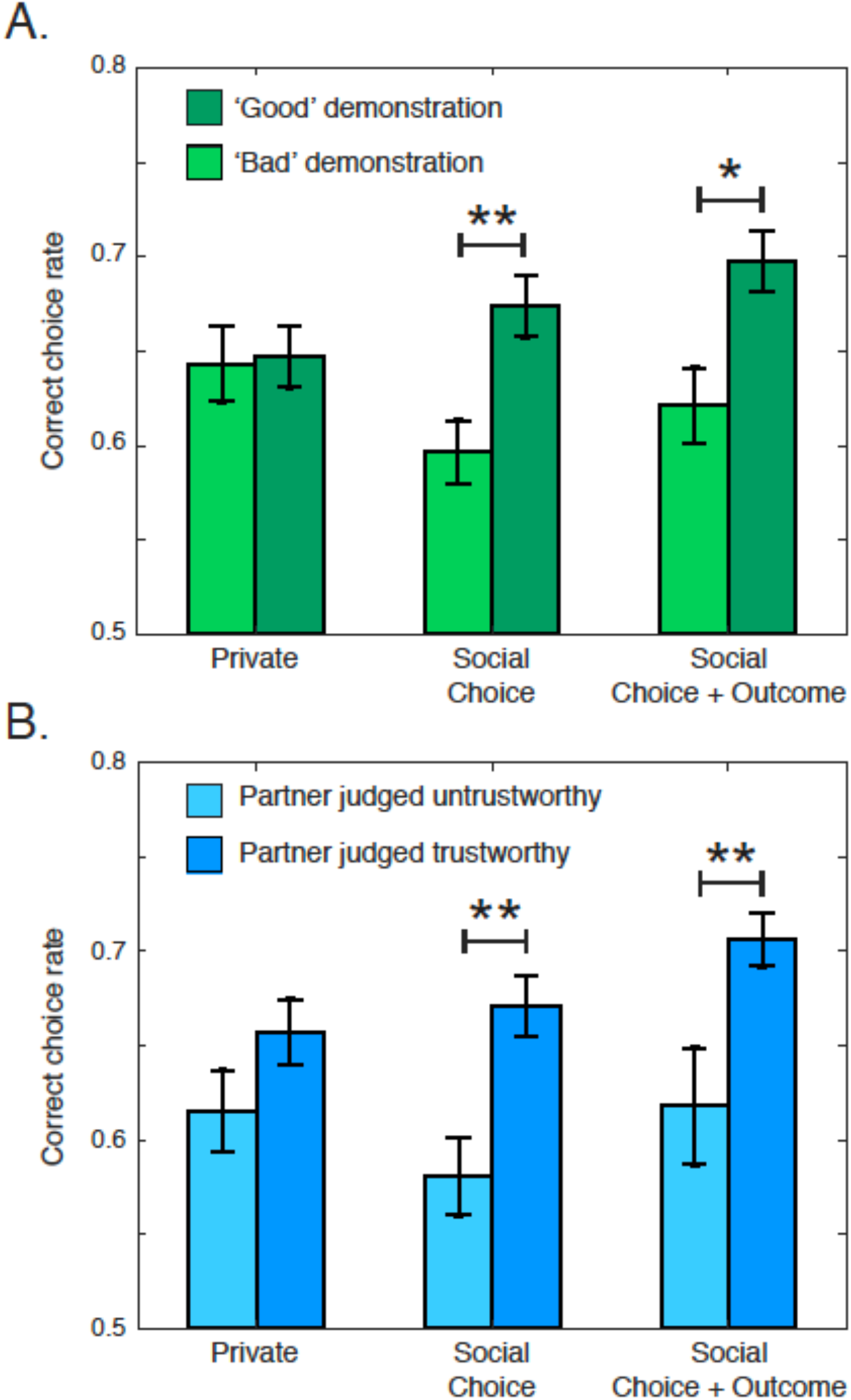
assessing social reinforcement learning. **(A) Effect of demonstrator’s behavior**. The demonstrator’s performance and the participant’s depression score influenced the correct choice rate in the ‘Choice’ and in the ‘Social-Choice+Outcome’ conditions, such that observing a virtual player with a low correct choice rate (‘Bad’ demonstration; light green) induced a lower correct choice compared to observing a virtual player with a high correct choice rate (‘Good’ demonstration; dark green) (‘Social-Choice’: *t*(198)= 3.17, *p* = .002; ‘Social-Choice+Outcome’: *t*(198)= 2.47, *p* = .014). **(B) Effect of perceived trustworthiness**. Participants who rated the demonstrator’s avatar as trustworthy had a higher correct choice rate in the ‘Social-Choice’ and ‘Social-Choice+Outcome’ conditions compared those who rated the avatar as untrustworthy (‘Social-Choice’: *t*(76)= 3.22, *p* =.002, ‘Social-Choice+Outcome’: *t*(76)= -2.87, *p* =.005)

In order to confirm that participants actually integrated the virtual demonstrator as a social partner, we measured the influence of participants’ rating of trustworthiness of the demonstrator’s face on social learning. An effect of perceived trustworthiness evaluations was found, such that participants who perceived the demonstrator’s avatar as more trustworthy had higher correct choice rates in the ‘Social-Choice’ (*t*(98) = 3.17, *p* = .002) and in the ‘Social-Choice+Outcome’ conditions (t(98) = 2.58, *p* = .012) but not in the ‘Private’ condition (t(98) = 1.08, *p* > .250; **Figure 2B**). This effect of the social evaluation of the demonstrator’s avatar confirms that participants processed the information in a social context.

### Correlation between depressive symptoms and performance

To test the effect of depression, the mixed linear logistic regression also included depressive symptoms as a between-subject variable. Importantly, anxiety, which is a comorbid trait of depression *(21, 22)*, was also included as a controlling factor (the regression also included a range of controls listed in **Table 2**). The analysis revealed a significant effect of depressive symptoms, such that the higher the depressive symptoms, the lower the rate of correct choices in the ‘Social-Choice’ condition compared to the ‘Private’ condition (*t*(489) = -2.64, *p* = .009; no other significant effect of depression and anxiety scores was evidenced: all *ps* > .250; **Figure 3A**). Importantly, the negative effect of depression in the ‘Social-Choice’ condition was particularly robust, because it was found in both the discovery and the replication sample and in the blocks with stable and reversal contingencies (within-subject) (**Figure 4**).

**Table 2:** statistical effect (mixed linear model) of social task conditions (‘Social-Choice’ and ‘Social-Choice+Outcome’), performance of the virtual demonstrator, perceived trustworthiness (‘Trustworthiness’), psychiatric scores (‘Depression’ and ‘Anxiety’), and their interaction when comparing to the ‘Private’ condition.

| Effect | Coefficient | SEM | DF | T-value | P-value |
| --- | --- | --- | --- | --- | --- |
| Social-Choice | -0.18 | 0.06 | 489 | -2.58 | 0.010* |
| Social-Choice+Outcome | -0.13 | 0.07 | 489 | -1.92 | 0.055° |
| Demonstrator performance | 0.01 | 0.06 | 489 | 0.15 | 0.883 |
| Trustworthiness | 0.03 | 0.03 | 96 | 1.04 | 0.302 |
| Depression | 0.01 | 0.02 | 96 | 0.47 | 0.638 |
| Anxiety | -0.01 | 0.02 | 96 | -0.46 | 0.643 |
| Social-Choice x Demonstrator performance | 0.21 | 0.08 | 489 | 2.54 | 0.011* |
| Social-Choice+Outcome x Demonstrator performance | 0.20 | 0.09 | 489 | 2.18 | 0.030* |
| Social-Choice x Trustworthiness | 0.05 | 0.03 | 489 | 1.48 | 0.140 |
| Social-Choice+Outcome x Trustworthiness | 0.04 | 0.03 | 489 | 1.22 | 0.222 |
| Social-Choice x Depression | -0.05 | 0.02 | 489 | -2.64 | 0.009** |
| Social-Choice+Outcome x Depression | -0.00 | 0.02 | 489 | -0.00 | 0.999 |
| Social-Choice x Anxiety | 0.01 | 0.02 | 489 | 0.72 | 0.475 |
| Social-Choice+Outcome x Anxiety | -0.01 | 0.02 | 489 | -0.51 | 0.614 |

**Table 3:** Estimated model parameters for the actual participants and for the simulated virtual demonstrators (mean ± 95% c.i.).

|  | Participants | Virtual demonstrators |
| --- | --- | --- |
| $\beta_{\text{private}}$ | $2.20 \pm 0.47$ | $9.54 \pm 0.49$ (real: 10) |
| $\alpha_{\text{private}}$ | $0.58 \pm 0.05$ | $0.52 \pm 0.02$ (real: 0.50) |
| $\beta_{\text{social}}$ | $1.83 \pm 0.34$ | |
| $\alpha_{\text{social}}$ | $0.60 \pm 0.06$ | |
| $\kappa$ | $0.13 \pm 0.02$ | |
| $\alpha_{\text{observation}}$ | $0.46 \pm 0.06$ | |

**Figure 3:**
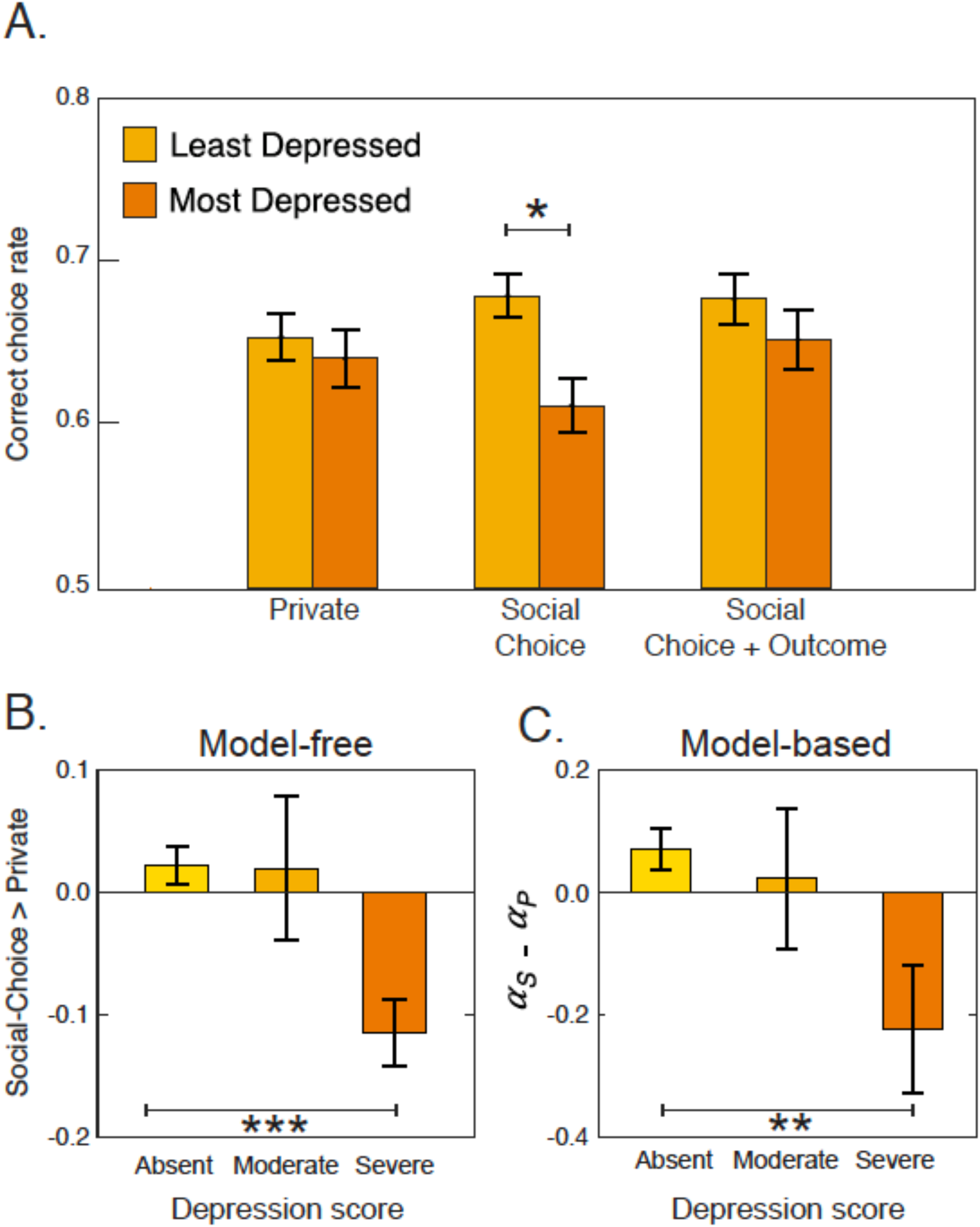
effect of depression on reinforcement learning in social contexts. **(A) Effect of depression**. The participant’s depression score influenced the correct choice rate in the ‘Social-Choice’ condition, such that participants with higher score had a lower correct choice rate in the ‘Social-Choice’ condition (median split: *t*(98) = -2.69, *p* = .008). **(B) Model-free classification**. The correct choice rate difference between the Private and the ‘Social-Choice’ conditions was significantly different between participants with ‘Absent’ and ‘Severe’ depressive symptoms (*t*(83) = 3.61, *p* < .001). **(C) Model-based classification**. The difference between the learning rate of the ‘Private’ and the social information contexts was significantly different between participants with ‘Absent’ and ‘Severe’ depressive symptoms (*t*(83) = -3.20, *p* =.002). Absent and Severe depressive symptoms correspond to scores < 8 and scores>10 on the HAD depression subscale, respectively.

**Figure 4:**
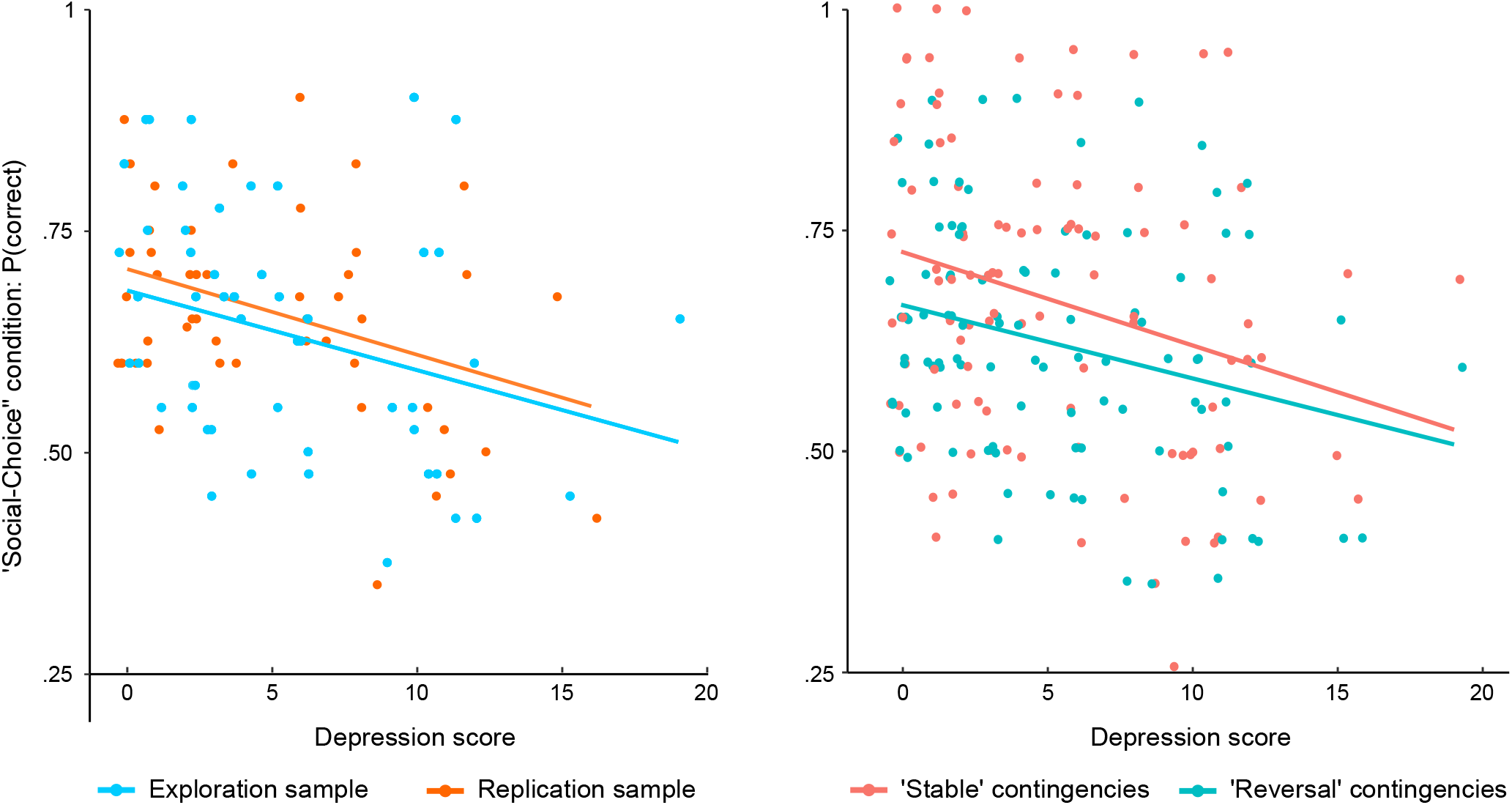
robustness of the result. The scatter plots represent the correlation between correct response rates in the ‘Social-Choice’ condition and depression scores separately for each experiment (right; exploration sample: *r* = -.29, *t*(48) = -2.15, *p* = .036; replication sample: *r* = -.37, *t*(48) = -2.75, *p* = .008) and reward contingency (right; exploration sample: *r* = -.27, *t*(98) = -2.83, *p* = .006; replication sample: *r* = -.27, *t*(48) = -2.81, *p* = .006; *r* and p: Pearson’s coefficient statistical significance.

Finally, we tested whether the difference in correct choice rates between the ‘Social-Choice’ and ‘Private’ conditions could accurately identify participants with ‘severe’ depression symptoms (i.e. scoring > 10 on the HAD depression subscale *(20)*). The classification analysis revealed that the performance difference between the ‘Social-Choice’ and the ‘Private’ condition identified the participants with ‘severe’ depressive symptoms with a good accuracy of 76 ± 1 % and both good a sensitivity, or True Positive Rate (78 ± 2%) and specificity, or True Negative Rate (63 ± 3%) of the classifier (**Figure 3B**).

### Computational model-based analyses

Although model-free analyses reveal a robust negative effect of depressive symptoms on learning in the ‘Social-Choice’ condition, they do not elucidate the cognitive mechanisms underlying this effect. Indeed, the effect of depressive symptoms could either be due to differences in social information processing, such as the demonstrator’s choices and outcomes (i.e. a *primary* social learning deficit) or to differences in the weighting of the information generated by participants’ own choices when social information is also available (i.e. a *secondary* social learning deficit or *audience effect).* These two hypotheses are hard to tease apart based on raw behavioral analyses, because both predict a reduced correct choice rate in the ‘Social’ conditions. Thus, to arbitrate between these two possibilities, we fitted a previously validated social reinforcement learning model *(14, 23)*. This model allows for biasing participants’ choice depending on their demonstrator’s choice in the ‘Social-Choice’ (i.e. *imitation*) condition and to update the value attributed to each symbol depending on the demonstrator’s outcome in the ‘Social-Choice+Outcome’ condition (i.e. *vicarious trial-and-error*). Compared to the original model and to directly assess the ‘socially induced individual learning deficit’ hypothesis *(14)*, we allowed participants to have different individual learning parameters in the ‘Private’ and in the two social conditions (‘Social-Choice’ and ‘Social-Choice+Outcome’ conditions; **Figure 5A**).

**Figure 5:**
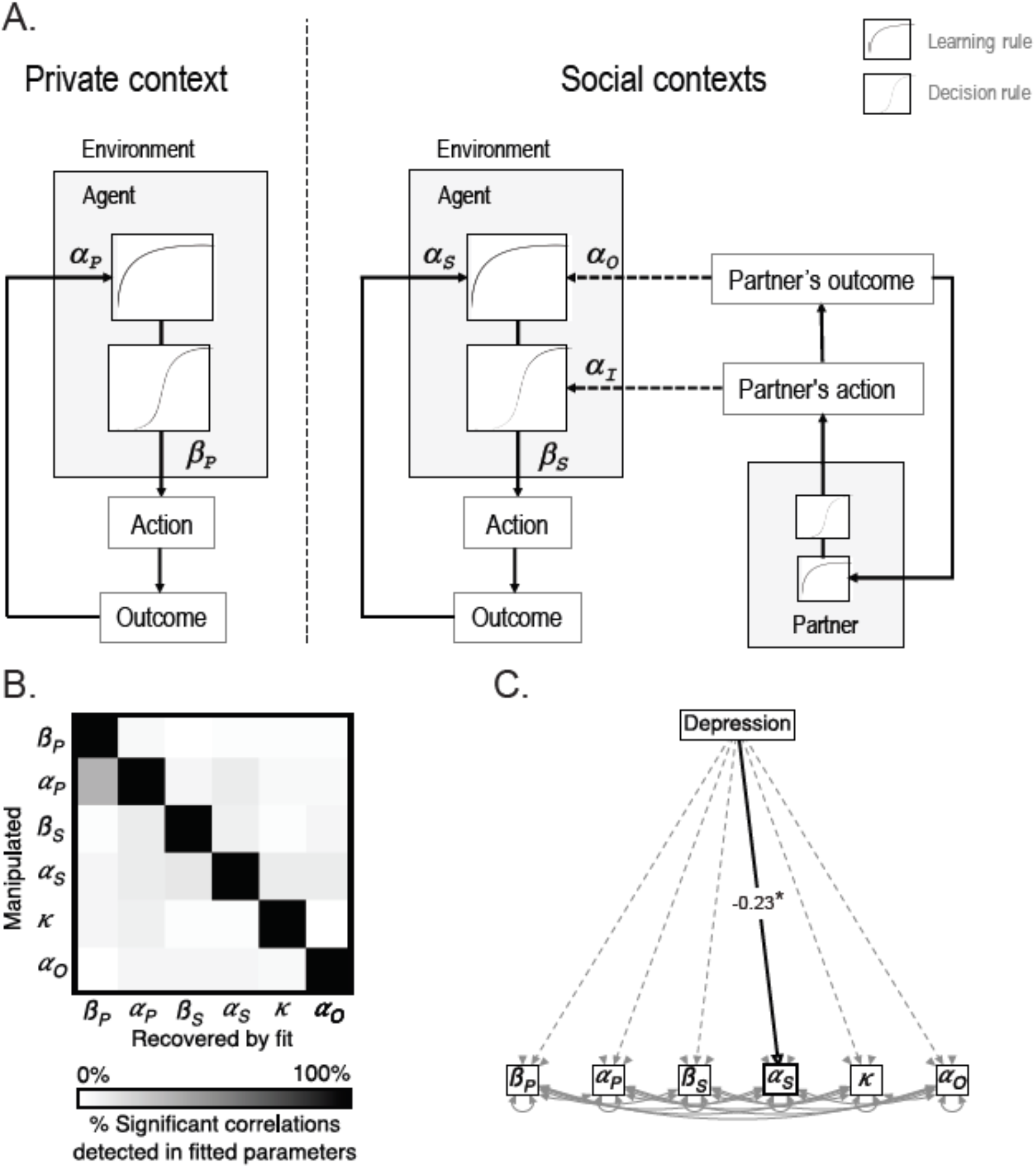
social reinforcement learning model and computational results. **(A) Computational model**. A social reinforcement learning model was fitted on participants’ behavior. In the ‘Private’ condition (‘Private context’), the model corresponded to a classical Q-learning (or Rescorla-Wagner) model. In Social context’ (‘Social-Choice’ and ‘Social-Choice+Outcome’ conditions), the model assumes that social information is integrated into the learning and decision process. Following Burke et al. (14), choice probability was updated based on the demonstrator’s action (imitation) in the ‘Social-Choice’ condition and the option value was updated when the demonstrator’s outcome was presented (counterfactual learning) in the ‘Social-Choice+Outcome’ condition. The proposed model also allows for different private parameters (learning rate, *α_S_*, and choice randomness, *β_S_)* being in the Social context. **(B) Parameter recovery**. To assess the sensitivity and the specificity of our model fitting procedure, we conducted a parameter recovery analysis. The matrix represents the percentage of significant correlations detected between different combinations of parameters. The diagonal cases correspond to the correlations that are accurately recovered; the other cases correspond to correlations that are spuriously recovered. **(C) Effect of depression on the model parameters**. Depression was specifically associated with a decrease in the private learning rate in the Social context α_S_), even controlling for the correlation between the different model parameters (structural equation modeling).

### Computational effects of depressive symptoms

We analyzed the model parameters fitted on participants’ actual behavior with structural equation modeling using depression scores as the independent variable. Higher depression scores were specifically associated with lower learning rates in the ‘Social’ conditions (depression: *z* = -2.41, *p* = .016; other *ps* > .199; **Figure 5B**). Interestingly, high depression scores were not solely associated with decreased learning rates in the ‘Social’ conditions, but also with decreased learning rates in the ‘Social’ conditions *compared* to the ‘Private’ condition (*t*(98) = -2.25, *p* = .027; **Figure 4C**), which indicates that the presence of social information decreased the learning rate of the most depressed participants. Adopting a computational psychiatry approach, we tested whether the difference in learning rates between the ‘Private’ and ‘Social’ conditions could identify ‘severe’ depressive symptoms (i.e. HAD depression subscale score above 10 (20)). The difference in learning rates detected participants with severe depressive symptoms with good accuracy (79 ± 1%), good sensitivity (84 ± 1%), but low specificity (53 ± 3%). A comparison between a classifier based on the model parameters and a classifier based on correct choice rates revealed that the model-based classifier was more accurate at detecting participants with ‘severe’ depressive symptoms (*t*(198) = 3.69, *p* < .001), and more sensitive (*t*(198) = 5.09, *p* < .001) but less specific (*t*(198) = -3.73, *p* < .001) than the classifier based on correct choice rates.

### Model simulations analyses

Model-based analyses indicated that depression severity specifically reduced individual learning rate in ‘Social’ conditions (*α_P_*): a parameter that is used both in the ‘Social-Choice’ and the ‘Social-Choice+Outcome’ condition. Model-free behavioral analyses showed that the learning deficit associated with depression severity was specific to the ‘Social-Choice’ condition. To ascertain that this computational result was compatible with our model-free observation, we ran the same statistical analysis on simulated data *(24)*. Crucially, data simulated using the fitted parameters accurately recovered the decrease in performance associated with depression scores in the ‘Social-Choice’ condition compared to the ‘Private’ condition (*t*(488) = -2.18, *p* = .030; Figure 4D) as well as the non significant effect of depression scores in the ‘Social-Choice+Outcome’ condition compared to the ‘Private’ condition (*t*(488) = -1.32, *p* = .188) and in the ‘Private’ condition (t(96) = -0.37, *p* > .250). Thus, the simulations captured the specificity of the behavioral effect of depression scores and illustrate that our model provides an accurate description of the data.

### Checking parameter recovery

As we were interested in the modulation of specific parameters by depression scores we tested whether our task allowed us to successfully retrieve a correlation between parameters in simulated datasets, an important quality check often referred to as ‘parameter recovery’ (24). To do so, we ran 100 sets of simulations for each parameter, each simulating 100 participants, with the parameter of interest correlating with an arbitrary variable and the other parameters being randomly fixed in a defined range. The simulated data were then fitted using our social reinforcement-learning model. Overall parameter recovery was very good, especially for the parameters of the social conditions, with significant correlations were found in the 100% of the simulated datasets (average correlation coefficient of the parameters: *r* = 0.73 ± 0.01). Importantly, the recovery of the correlations was specific to the manipulated parameter with false alarms detected in less than 10% of the cases except for learning rate and choice temperature in the ‘Private’ condition (which was not our condition of interest) (**Figure 5C**). This result indicates that it is very unlikely that a correlation of one of our parameters with participants’ HAD depression scores is actually due to an effect of depression scores on another parameter.

### Discussion

In the present study we assessed reinforcement learning with a behavioral paradigm involving both private and social contexts, while concomitantly assessing depressive and anxiety symptoms. First, we replicate previous findings showing that subjects integrate the demonstrator’s choices and outcomes, which is consistent with the idea that social learning processes (both in terms of imitation and vicarious trial-and-error) play a role in human reinforcement learning *(14, 15, 25–27)*. Second, we show that the severity of depressive symptoms is associated with a learning impairment that is specific to the learning context where participants are informed about the demonstrator’s choices (social context). This negative effect was robust to the inclusion of anxiety, and robust across experiments and outcome contingencies. Finally, computational analyses allowed us to characterize the effect of depressive symptoms as a secondary social learning deficit, i.e. a reduction of the learning rate in social contexts.

We found that depressive symptoms had a specific effect on imitation in the ‘Social-Choice’ condition. Crucially, the effect was robust to the inclusion of anxiety, which did not modulate performance in our task. That anxiety has no effect may come as a surprise given that previous studies have found that anxiety is associated with deficits in social and non-social reinforcement learning *(28)*. One possible explanation is that this anxiety is more strongly linked to classical fear conditioning, rather than reward-based instrumental learning *(29)*. Depressive symptoms might thus undermine social reinforcement learning in instrumental and reward-maximization contexts, while anxiety might affect the same processes when outcomes are independent from the participants’ choices (i.e. Pavlovian learning) and when outcomes have a negative valence (aversive contexts).

Model-free analyses *per se* do not allow us to pinpoint the psychological mechanisms underlying the negative effect of depressive scores on correct choice rates in the ‘Social-Choice’ context. The absence of interaction between the demonstrator’s performance and depressive symptoms suggests that depressive symptoms did not lead participants to disproportionally follow ‘bad examples’ or to be insensitive to ‘good examples’. However, interpretations based on negative results are, at best, unsafe. To formally characterize the psychological mechanisms of the detrimental effects of depressive symptoms we thus turned to model-based analyses.

We fitted subjects’ choice with a slightly modified version of a previously validated social reinforcement-learning model (14). As in standard algorithms, the model assumes that subjects learn option values via the calculation of a reward prediction error, which are moderated by a learning rate (**α**_*P*_) and that choices are generated via a soft-maximization process, whose stochasticity is governed by a temperature (**β**_*P*_) *(30).* In addition to this ‘private’ learning module, the model also displays sensitivity to social information: in the ‘Social-Choice’ condition the demonstrator’s choice biases the subsequent subject’s choice (the magnitude of this effect is governed by an imitation rate **κ**) and in the ‘Social-Choice+Outcome’ condition the demonstrator’s outcome is integrated into the subject’s value function with a vicarious learning rate (**α**_0_). Finally, we also allowed for different private learning rates and temperatures in the ‘Social’ contexts (**α**_S_ and **β**_S_). This precise model parameterization allowed us to disentangle two different hypotheses concerning the drop in performance associated with depressive symptoms in the ‘Social-Choice’ condition. A correlation between depression scores and imitation rates and/or vicarious learning rates would imply what we define a ‘primary’ social learning impairment (i.e. an impairment of the social learning processes *per se*). On the contrary, a correlation between the ‘Social’ context-specific learning rate and/or temperature would imply a ‘secondary’ social learning impairment (i.e. an impairment of the private learning processes in presence of social information). We found that depressive scores negatively correlated with the private learning rate in the social context (**α**_*S*_), thus indicating that the effect is consistent with a secondary impairment and is specific to the learning (as opposed to the decision) process. In other words, our computational results suggest that one possible way in which depressive symptoms affect learning in social contexts is conceptually similar to a negative audience effect *(31, 32)*, where the presence of social signals (the demonstrator’s choices) induces a reduction of subjects’ instrumental performance.

From a methodological point of view, our study exemplifies how computational approaches can provide new insights on the way in which cognitive processes vary with clinical symptoms. Indeed, computational modeling demonstrated that the effect of depressive symptoms was selective of the way individual information was processed *(33, 34)*. It is worth noting that these conclusions were only allowed after a careful testing of the ability of our task to precisely identify which model parameter would be influenced by depressive symptoms *(24)*. The exact cognitive and psychological mechanisms that mediate the negative effect of social signals in instrumental performance remain to be characterized. One possibility given that depression is associated with lower cognitive functioning in general *(35)* is that the mere presence of others exacerbates these difficulties by capturing already scarcer attentional resources. Alternatively, negative perception of self and negative comparison to others are core symptoms of depression *(36)*. Therefore, it is possible that the most depressed participants perceived their demonstrator’s behavior as more reliable, thus underweighting the information they acquired through their own experience.

Our results provide new evidence that depression-related reward learning deficits are highly context-dependent *(3–5)*, and suggest that the differences in learning rates associated with depressive symptoms may only arise in social contexts *(5, 9)*. Crucially, our results suggest that supposedly neutral aspects of the experimental setup (such as whether or not the task is done in the presence or absence of an experimenter), may affect the results and explain inconsistent findings *(42)*. In line with recent propositions, our results also suggest that a deeper investigation of socio-cognitive impairments in depression may provide important new insights *(10, 11)*. Finally, we suggest that developing tools assessing reward learning outside and inside social contexts (characterized either by the presence of another player or by the social nature of the outcomes *(43)*) may prove useful to improve diagnosis and personalize treatments of depressive syndromes in the long term.

An obvious limitation of our study, is that we did not control for participants’ actual diagnosis and treatment, which may be problematic since medication interacts with decision-making in depression *(44)*. Therefore, our results would benefit from being replicated in carefully controlled population, while controlling for medication status and medical history. This replication would allow us to further measure the diagnostic value of our behavioral task and associated computational model-based analyses. Indeed, in the present study, we only tested its ability to detect high depressive scores as identified by a self-rated scale *(20)*. It would be particularly interesting to test whether our behavioral and computational measures improve existing self-assessments that detect clinically diagnosed cases of depression.

Our results have implications beyond their clinical relevance. Consistent with the ‘social learning theory’ participants imitated demonstrators’ choices (‘Social-Choice’ condition) and learned from their outcomes (‘Social-Choice+Outcome’ condition) *(13, 14)*. At the behavioral level, these two psychological processes were manifest in the fact participants’ performance was modulated by the demonstrators’ performance. In particular, we found that participants observing a demonstrator performing ‘well’ performed better in the social compared to the private learning context. Importantly the opposite was also true: participants observing low performing demonstrators displayed lower performance in the social compared to the private context. This latter result is in apparent contrast with the normative view that imitation should be biased toward successful individuals in order to be evolutionary adaptive *(45–47)*. This is also in contrast with recent empirical evidence using a very similar paradigm and showing that imitation rate is modulated by the actual performance of the demonstrator, so that demonstrators making random (i.e., non reward-maximizing) decisions are less imitated *(15)*. Two differences between the previous design and ours may explain this discrepancy. First, the previous study involved mild electric shocks (primary reinforcer), while our study involved abstract points to be converted into money (secondary reinforcer). More importantly perhaps, the previous design involved a between-subjects design with two groups of participants paired either with a consistently good or with a consistently bad participant, while in our experiments the performance of the demonstrator was allowed to fluctuate in a within-subject manner around an optimal behavior. Therefore, it could also be argued that our experiment is not well-suited for measuring demonstrators’ performance effects on participants’ imitation behavior as such effects require a relatively long and stable reputation building process *(48, 49)*.

The question remains whether or not in our task social learning (imitation and vicarious trial-and-error) engaged domain-specific social cognitive module or domain-general information processing modules. In the absence of additional data (such as neuroimaging) we cannot provide a definitive answer. However, evidence from post-learning face ratings provides some clues *(50)*. We found a positive correlation between performance in the social contexts and the demonstrator’s judgment of trustworthiness. Even if we cannot infer a causal link and its direction from the post-learning face evaluation, these results suggest that a specific socio-cognitive module (face evaluation) correlated with instrumental performance, thus demonstrating the engagement of social information-specific processing and our reinforcement learning task.

## Materials and Methods

### Participants

Two independent cohorts of 100 American participants, similar in terms of reported age (mean reported age across the two cohorts: 33.39 ± 2.03) and of reported male/female ratio (mean reported male/female ratio across the two cohorts: 35%; see Table) were recruited via Amazon Mechanical Turk to participate in this online study. Each participant received a fixed 4$ amount for completing the 40-minute task to which a bonus earned during the experiment was then added (average bonus: 0.49$). Participant received a description of the study and signed an informed consent before starting the experiment. The study was approved by the ethical committee. The first cohort corresponded to a ‘discovery experiment’ where we explored the relation between instrumental performance and clinical scores; the second cohort corresponded to a ‘replication experiment’ where we tested the robustness and replicability of the effect identified in the first experiment.

### Experimental design

Participants performed the probabilistic instrumental learning task described in the Results section (**Figure 1A**). The task was programmed on Qualtrics and was composed of six learning blocks of 20 trials each. In each block, participants had to choose between two cues. Cues were characters of the agathodaimon font and were always presented in pair and only in one block *per* subject. The cue-to-condition attribution was randomized across subjects. Participants made their choice by pressing the E or P keys to choose the leftmost or rightmost symbol. Participants were given no explicit information on reward probabilities, which they had to learn through trial and error. In addition, they were encouraged to accumulate as many points as possible, with their final amount of points being translated into bonus money at the end of the experiment (conversion rate: 40 points equals 1$ bonus). In each pair, cues were associated with reciprocal reward probabilities (20/80% or 30/70%). For instance, in a 30/70% pair, the most rewarded cue provided a positive outcome (+1 point) 70% of the times and a negative outcome (-1 point) 30% of the time, while the less rewarded cue provided a negative outcome 70% of the time and a positive outcome 30% of the time.

Participants were told they had been paired with another player at the beginning of the experiment with whom they played in turn in each trial. As in previous studies (Suzuki et al. Neuron 2012), the behavior of the demonstrators was determined by a reinforcement learning algorithm (Q-learning) with a reasonable set of free parameters (*α* = 0.5, *β* = 10; see below for a description for the Q-learning and its parameters). To avoid social perceptual biases, the other player was represented by a neutral avatar, chosen to be generally perceived as neither dominant or submissive nor trustworthy or untrustworthy (51). Participants had to choose their own avatars in a set of other 16 identities (8 female, 8 male) at the beginning of the task. Participants performed this task in three different contexts with different amounts of social information: a ‘Private’ condition in which they did not have access to the demonstrator’s behavior, a ‘Social-Choice’ condition in which participants could see the demonstrator’s behavior but not their outcomes and a ‘Social-Choice+Observation’ in which participants could observe the demonstrator’s decisions and outcomes. Importantly, participants performed each condition (‘Private’, ‘Social-Choice’ and ‘Social-Choice+Outcome’) twice. In the ‘Stable’ type of contingency, outcome probabilities were set at 30/70% and did not change during the block. In the ‘Reversal’ type of contingency, outcome probabilities were set at 20/80%.and was inverted across cue after 10 trials (in average). Finally, at the end of the experiment, participants rated their demonstrator’s avatar on three personality traits (trustworthiness, dominance and competence) and completed the Hospital Anxiety and Depression Scale *(20)* as well as the Peters et al. Delusions Inventory, that was included in the exploratory analysis of the ‘discovery’ experiment and then discarded in absence of any significant effect and its inclusion did not affect the effect of depression.

### Statistical analyses

#### Percentage of correct choices

Percentage of correct choices were extracted for each block and used as a dependent variable. A mixed linear regression with both random intercept and random slopes was conducted on correct choice rates taking participants’ ID as a random factor, condition (‘Private’, ‘Social-Choice vs ‘Social-Choice+Outcome’) as within-subject variables and depression and anxiety scores as well as demonstrator’s performance and trustworthiness judgment as between-subject variables (**Table 2**).

#### Diagnostic value

Out of sample tests were used to assess the diagnostic value of our task, i.e., its ability to distinguish participants scoring below the ‘severe symptoms’ threshold in depression scale from those above this threshold. 50 participants were randomly extracted from the entire sample and used to optimize a classifier of ‘severe’ depressive symptoms (HAD depressio subscale score above 10 *(20)*) using either the difference in correct choice rates between the ‘Social-Choice’ and the ‘Private’ conditions (model-free measure) or the difference in learnin¢ rates between the Private and social information conditions (*α_S_* minus *α_P_*; model-based measure; see below). The classifier and the associated optimal cut-off was tested on the 50 remaining participants. This operation was repeated 100 times in order to estimate the average accuracy, sensitivity and sensibility of the classifiers.

### Computational analyses

#### Computational model

To fit the behavioral data, we used a social reinforcement learning model previously presentee by Burke et al. *(14)*. Individual learning and decision-making where modeled with classica softmax (eq.1) and delta-rule (eq.2) functions, respectively governed by learning rate and choice randomness (or temperature) parameters:

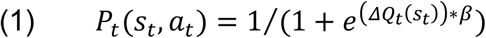

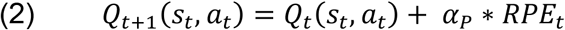

Where *RPE_t_* is the reward prediction error calculated as follows (eq.3):

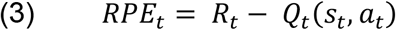

The only change we made was the inclusion of different learning rates and inverse temperature parameters in the ‘Private’ (*α_P_, *β*_P_*) and social information (*α_S_, *β*_S_*) conditions. During the ‘Social-Choice’ condition, the model assumes that the Demonstrator’s choice induces an ‘action’ prediction error (*APE_t_*; (eq.4)), which measures how surprising the Demonstrator’s choice is, given the subject’s current estimate of the option values:

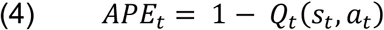

The *APE_t_* is then used to bias the subject’s choice probability (eq.5) in the subsequent trial and the effect is scaled by a parameter *κ* ∈ {0-1}:

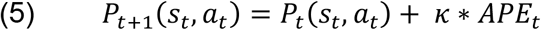

Finally, in the ‘Social-Choice+Outcome’ trials, the model assumes that the Demonstrator’s outcome induces an ‘observational’ reward prediction error (eq.5), which is scaled by observational learning rate *α*_0_ ∈ {0-1} (eq.6):

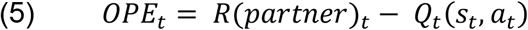

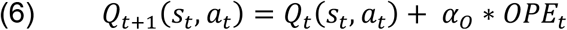

To sum up, our computational modeling allowed us to address both *primary* social learning deficits *(i.e.* learning deficits captured by the parameters *κ* and *α*_0_, which are specific to social information) and *secondary* social learning deficits *(i.e.* learning deficits captured by the parameters *β_S_* and *α_S_*, which are specific to individual learning in contexts where social information is available).

We optimized the model parameters by minimizing the Laplace approximation to the model evidence (log of the posterior probability: LPP) (eq.7):

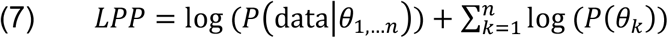

Where D represents the data, *θ*_1,…*n*_ the model, and *θ_k_* represents one of the *n* parameters of the computational model. The LPP represents a trade-off between the model’s accuracy and complexity: it increases with the likelihood of the model given the data (a measure of fit) and decreases with the number of parameters. By including priors over the parameters, this method avoids degenerate parameter estimation. In our analysis, the priors were defined as a gamma function (gampdf(1.2,5)) for the temperature parameters (range: 0<β<Infinite) and as a beta function (betapdf(1.1,1.1)) for the learning and imitation rates (ranges: 0<α<1, 0<κ<1) as described in *(52)*

Importantly, LPP analysis suggested that the social reinforcement learning fit the data better than a simple Q-learning model without social influence, even accounting for its extra-complexity (social reinforcement learning model: posterior probability: 90 ± 3 %; exceedance probability: 100%). As a control analysis, in order to ensure that our model comparison criterion was not over-fitting prone, we fit the behavior of the virtual demonstrators that we generated with a Q-learning model. This model recovery analysis *(24)* correctly indicated that the simple Q-learning model explained the demonstrators’ data better (social reinforcement learning model: posterior probability: 100 ± 0 %; exceedance probability: 100%).

Because the model parameters were correlated with each other (maximal correlation: *r* = 0.53), we used structural equation modeling to analyze the influence of depression scores on the model parameters. This technique allowed to test the influence of depression scores on each parameter while simultaneously accounting for the inter-correlations of the dependent variables (the model free parameters) and of the independent variable (the depression score).

#### Model simulation analyses

Finally, we assessed the ability of the model to recover the observed behavioral effect of depressive symptoms using model simulations (24). For each participant, we simulated behavioral data for each condition based on their best fitting parameters. Importantly, a simulated demonstrator was also generated, such that the simulated data were completely independent of the contingencies actually experienced by the participants. This procedure was repeated 100 times, to avoid any effect of participant’s and demonstrator’s history of choice and outcomes. Analysis of the recovered percentage of correct choices was ran on the averaged rates of correct choices across the 100 simulations using a linear mixed regression taking the exact same predictors as the mixed general linear model used for analyzing participants’ percentage of correct choices.

## H2: Acknowledgments

SP is supported by an ATIP-Avenir grant (R16069JS) Collaborative Research in Computational Neuroscience ANR-NSF grant (ANR-16-NEUC-0004), the Programme Emergence(s) de la Ville de Paris and the Fyssen foundation. LS was supported by a PHD fellowship of the ENS/PSL. The Institut d’Etude de la Cognition is supported financially by the LabEx IEC (ANR-10-LABX-0087 IEC) and the IDEX PSL* (ANR-10-IDEX-0001-02 PSL*). The funders had no role in the conceptualization, design, data collection, analysis, decision to publish, or preparation of the manuscript.

